# Forecasting type-specific seasonal influenza after 26 weeks in the United States using influenza activities in other countries

**DOI:** 10.1101/705855

**Authors:** S. B. Choi, J. Kim, I. Ahn

**Affiliations:** Department of Data–centric Problem Solving Research, Korea Institute of Science and Technology Information, Daejeon, Republic of Korea; Center for Convergent Research of Emerging Virus Infection, Korea Research Institute of Chemical Technology, Daejeon, Republic of Korea

**Keywords:** Cross–correlation, FluNet, Influenza, Forecast, Deep learning

## Abstract

To identify countries that have seasonal patterns similar to the time series of influenza surveillance data in the United States and other countries, and to forecast the 2018–2019 seasonal influenza outbreak in the U.S. using linear regression, auto regressive integrated moving average, and deep learning. We collected the surveillance data of 164 countries from 2010 to 2018 using the FluNet database. Data for influenza-like illness (ILI) in the U.S. were collected from the Fluview database. This cross-correlation study identified the time lag between the two time-series. Deep learning was performed to forecast ILI, total influenza, A, and B viruses after 26 weeks in the U.S. The seasonal influenza patterns in Australia and Chile showed a high correlation with those of the U.S. 22 weeks and 28 weeks earlier, respectively. The *R*^2^ score of DNN models for ILI for validation set in 2015–2019 was 0.722 despite how hard it is to forecast 26 weeks ahead. Our prediction models forecast that the ILI for the U.S. in 2018–2019 may be later and less severe than those in 2017–2018, judging from the influenza activity for Australia and Chile in 2018. It allows to estimate peak timing, peak intensity, and type-specific influenza activities for next season at 40^th^ week. The correlation for seasonal influenza among Australia, Chile, and the U.S. could be used to decide on influenza vaccine strategy six months ahead in the U.S.

## Introduction

Seasonal influenza viruses are a significant public-health problem that causes a great many deaths worldwide every year [1]. The annual recurrence of seasonal epidemics is attributed to the continued evolution of seasonal influenza viruses, which enables them to escape the immunity that is induced by prior infections or vaccinations, as well as to the ability of those viruses to be transmitted efficiently human‑to‑human via respiratory droplets, direct contact, and fomites [1]. Currently, the human influenza A subtypes H1N1 and H3N2, as well as influenza B viruses, are the most commonly encountered variants worldwide [2]. Vaccination for seasonal influenza is the primary tool for reducing its morbidity and death rate [3]. If public-health officials could accurately predict where and when epidemics would take off, they could allocate resources and implement preventative measures to mitigate morbidity and the death rate, or save resources in epidemic situations expected to diminish on their own [4].

Early prediction of the international spread of viruses during a potential pandemic can guide public-health actions globally [5]. Several previous studies have focused on predicting the incidence rate, peak time, or onset time of influenza-like illness (ILI) using data from online volunteer participants, ILI-related queries on Google, Wikipedia logs, or a combination of several data sources, including temperature and humidity [6–8]. However, the previous studies have focused on short-term forecasts for up to four weeks, and few studies have predicted seasonal influenza epidemics by using surveillance data from other countries.

Because influenza epidemics are acute, the long-term circulation of influenza viruses in the human population is driven by the global movement of viruses. The extent to which viruses move internationally versus persisting locally during different epidemics has been of interest since at least the 1800s [1]. High-quality influenza surveillance systems are needed to enable countries to better understand influenza epidemiology, including disease incidence and severity, and help them implement appropriate prevention strategies [9].

To identify countries with seasonal patterns and influenza outbreaks that are similar to but precede those of the United States, we used FluNet surveillance data to investigate cross– correlation between the U.S. and other countries, in terms of time-series of the ILI, total influenza (Total INF), influenza A (INF A), and influenza B (INF B) viruses. Our hypothesis was that similar seasonal patterns of influenza outbreaks between two countries over the years are associated with influenza activity in the following year. Knowing about such an association may help clinicians to predict the pattern of influenza incidence in the next season. The prediction model allows to estimate peak timing, peak intensity, and type-specific influenza activities for next season at 40^th^ week.

## Methods

### Data collection

Data for ILI in the U.S. were collected from the Fluview database of the Centers for Disease Control and Prevention in the U.S. The agency’s website (https://gis.cdc.gov/grasp/fluview/fluportaldashboard.html) provides both new and historical data. The CDC’s ILI is freely distributed and available through ILInet [10]. From this website, we can obtain the CDC’s dataset on weighted ILI (%).

Influenza surveillance data were collected from the FluNet database of the WHO Global Influenza Surveillance Network (URL: http://apps.who.int/flumart/Default?ReportNo=12) [3, 11]. These data are uploaded to the FluNet database every week by the countries in the network [3]. The FluNet database contains the following variables, reported by 164 countries: influenza transmission zone, number of specimens, number of INF A and INF B detected by subtype, and number of influenza-positive viruses. We collected the surveillance data of the 164 countries from the 40^th^ week in 2010 to the 40^th^ week in 2018. We set the starting point for our research data in 2010, because influenza season during 2009 was an atypical season, with the introduction of a novel pandemic strain (INF A H1N1 pdm09) [12]. Missing data were replaced by a zero because some countries conducted surveillance only during weeks with high influenza activity. Total INF, INF A, and INF B were defined as the number of influenza subtypes detected among processed specimens. The overall flow chart for this study is presented in Fig 1.

**Fig 1.**
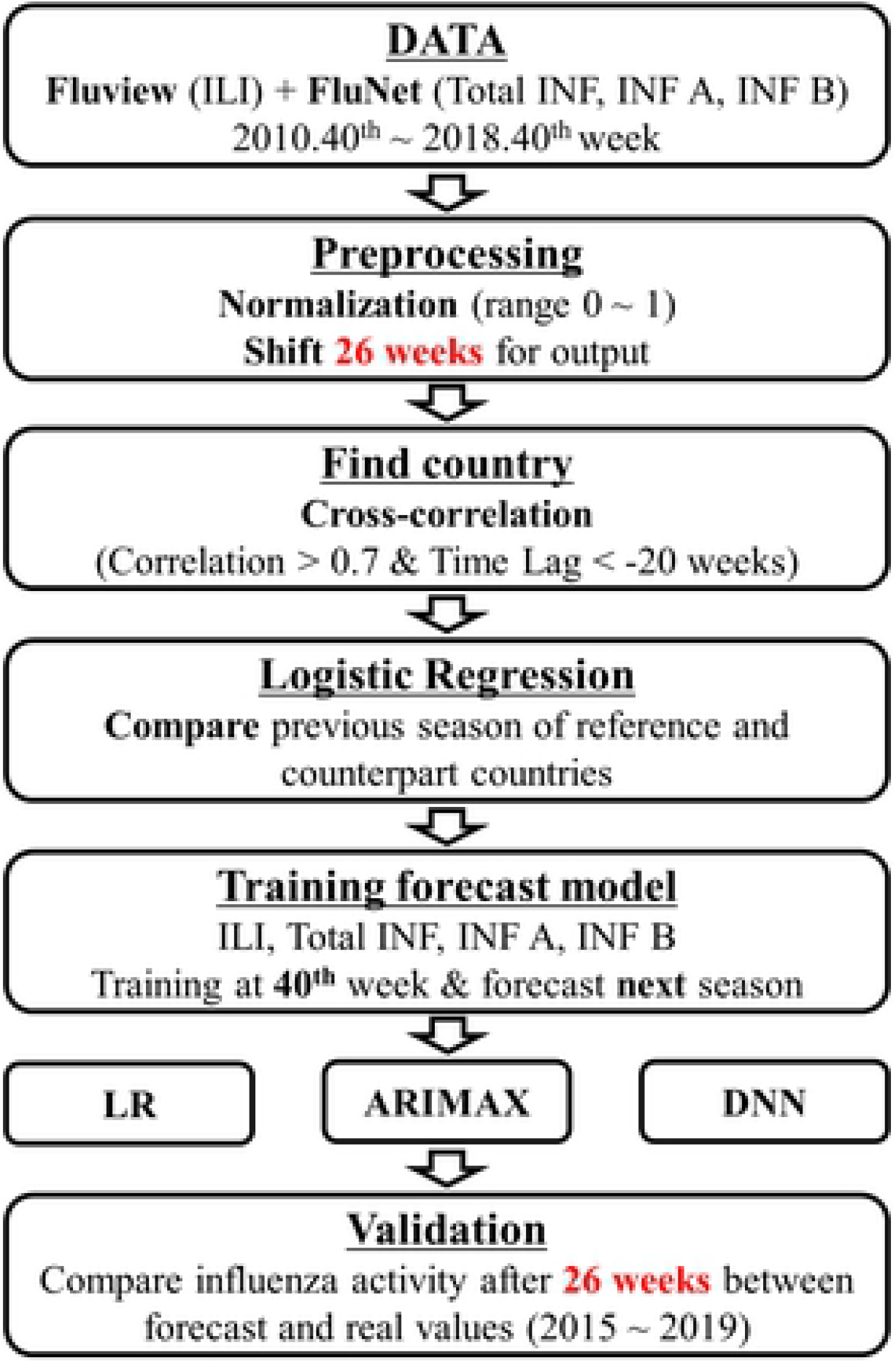
The flow chart for this study.

### Statistical analysis

All surveillance data were normalized by country, and each value was changed into a range of variables from “0” to “1” using the following equation: {*Z*_i_ = [*x*_i_ − min (*x*)]/[max (*x*) − min (*x*)]}, *x*_i_ was the value of i^th^ weeks for each country. In two time periods, cross-correlations were analyzed using Pearson’s correlation, with a time-lag range of ± 30 weeks. Cross-correlation allows the time lag between two time series to be identified [13]. If the blue waveform of the reference country correlates with the green waveform of country A with a time lag of −2 weeks, the peak or onset of the reference country can be identified as occurring 2 weeks later than that of country A (S1 Fig). Time-series for ILI, Total INF, INF A, and INF B in the U.S. as reference were analyzed with the Total INF, Total INF, INF A, and INF B, respectively, in the comparison country using cross-correlation. For example, ILI in the U.S. was compared to Total INF in each comparison country. We selected counties with a time lag of −20 weeks or less and a correlation coefficient of 0.7 or more in Table 1.

**Table 1.** Maximum correlation coefficient and time lag with time series of influenza surveillance data in the United States and other countries (Top 16)

|  | ILI |  |  | Total INF |  |  | INF A |  |  | INF B |  |  |
| --- | --- | --- | --- | --- | --- | --- | --- | --- | --- | --- | --- | --- |
|  | Country | Corr | Time lag | Country | Corr | Time lag | Country | Corr | Time lag | Country | Corr | Time lag |
| 1 | Canada | 0.872 | 1 | Australia* | 0.892 | -22 | Australia* | 0.896 | -22 | Norway | 0.884 | 0 |
| 2 | Australia* | 0.855 | -22 | Canada | 0.885 | 1 | Canada | 0.829 | 0 | Sweden | 0.864 | 1 |
| 3 | Chile* | 0.784 | -28 | UK | 0.825 | 0 | Chile* | 0.802 | -28 | UK | 0.855 | -2 |
| 4 | Germany | 0.781 | 4 | Norway | 0.803 | 2 | UK | 0.747 | 0 | Croatia | 0.854 | -2 |
| 5 | Norway | 0.766 | 2 | Denmark | 0.796 | 4 | Laos | 0.656 | -19 | Canada | 0.851 | 0 |
| 6 | Sweden | 0.751 | 4 | Chile* | 0.777 | -29 | Bangladesh | 0.656 | -30 | Denmark | 0.848 | 1 |
| 7 | Iceland | 0.744 | 3 | Spain | 0.775 | -2 | Luxembourg | 0.650 | 4 | Switzerland | 0.841 | -4 |
| 8 | Japan | 0.731 | -1 | Iceland | 0.770 | 2 | China | 0.639 | 0 | Ireland | 0.840 | -3 |
| 9 | Croatia | 0.719 | 1 | Sweden | 0.766 | 3 | Nepal | 0.631 | -22 | Austria | 0.830 | -1 |
| 10 | UK | 0.706 | 1 | Malta | 0.762 | -2 | Oman | 0.612 | -11 | Spain | 0.821 | -4 |
| 11 | Latvia | 0.705 | 5 | Switzerland | 0.758 | -1 | Denmark | 0.611 | 7 | Serbia | 0.821 | -1 |
| 12 | Austria | 0.704 | 3 | China | 0.750 | -1 | Spain | 0.611 | 0 | Lithuania | 0.804 | -1 |
| 13 | Spain | 0.694 | -1 | Austria | 0.733 | 2 | France | 0.605 | -1 | Qatar | 0.803 | -8 |
| 14 | Denmark | 0.693 | 5 | Portugal | 0.725 | 0 | Colombia | 0.600 | 20 | Portugal | 0.797 | -3 |
| 15 | Luxembourg | 0.685 | 3 | Germany | 0.714 | 3 | Iceland | 0.582 | 2 | Estonia | 0.794 | 1 |
| 16 | Czechia | 0.684 | 3 | Ireland | 0.709 | -1 | Algeria | 0.575 | 1 | Australia* | 0.792 | -25 |

Corr, maximum correlation coefficient; INF, Influenza; ILI, Influenza-like illness; UK, United Kingdom of Great Britain and Northern Ireland; US, United States of America
\* Countries with a correlation coefficient of 0.7 or more and -20 or less week time lag from 2010 to 2018

Linear regression analysis (LR) was used to evaluate the relationship between influenza surveillances in the U.S. after 26 weeks and those in selected countries from the 40^th^ week in 2010 to the 40^th^ week in 2018. LR 1 used influenza data from the U.S. after 26 weeks as dependent variable and previous seasonal data from the U.S. as independent variable. LR 2 used influenza activities from selected countries, and LR 3 used both previous seasonal data from the U.S. and influenza activities from selected countries data in Table 2.

**Table 2.** Linear regression analysis for influenza surveillance in selected countries with time lag and those of previous season in the U.S. from the 40^th^ week in 2010 to the 40^th^ week in 2018.

| ILI for U.S. after 26 week – Dependent Variable |  |  |  |  |
| --- | --- | --- | --- | --- |
| Model | Independent Variables | Adj. R-squared | Beta [95% CI] | P-value |
| LR 1 | ILI - U.S. (before 26 week) | 0.537 | 0.857 [0.778, 0.936] | < 0.001 |
| LR 2 | Total INF - Australia (present) | 0.761 | 0.645 [0.586, 0.703] | < 0.001 |
|  | Total INF - Chile (present) |  | 0.495 [0.439, 0.551] | < 0.001 |
| LR 3 | ILI - U.S. (before 26 week) | 0.809 | 0.338 [0.271, 0.405] | < 0.001 |
|  | Total INF - Australia (present) |  | 0.514 [0.455, 0.572] | < 0.001 |
|  | Total INF - Chile (present) |  | 0.380 [0.326, 0.435] | < 0.001 |
| Total INF for U.S. after 26 week – Dependent Variable |  |  |  |  |
| LR 1 | Total INF - U.S. (before 26 week) | 0.475 | 1.076 [0.964, 1.188] | < 0.001 |
| LR 2 | Total INF - Australia (present) | 0.729 | 0.669 [0.612, 0.727] | < 0.001 |
|  | Total INF - Chile (present) |  | 0.348 [0.293, 0.403] | < 0.001 |
| LR 3 | Total INF - U.S. (before 26 week) | 0.741 | 0.256 [0.141, 0.372] | < 0.001 |
|  | Total INF - Australia (present) |  | 0.559 [0.484, 0.634] | < 0.001 |
|  | Total INF - Chile (present) |  | 0.322 [0.267, 0.376] | < 0.001 |
| INF A for U.S. after 26 week – Dependent Variable |  |  |  |  |
| LR 1 | INF A - U.S. (before 26 week) | 0.387 | 0.900 [0.788, 1.012] | < 0.001 |
| LR 2 | INF A - Australia (present) | 0.690 | 0.577 [0.519, 0.636] | < 0.001 |
|  | INF A - Chile (present) |  | 0.471 [0.409, 0.532] | < 0.001 |
| LR 3 | INF A - U.S. (before 26 week) | 0.708 | 0.254 [0.152, 0.355] | < 0.001 |
|  | INF A - Australia (present) |  | 0.479 [0.410, 0.548] | < 0.001 |
|  | INF A - Chile (present) |  | 0.434 [0.373, 0.495] | < 0.001 |
| INF B for U.S. after 26 week – Dependent Variable |  |  |  |  |
| LR 1 | INF B - U.S. (before 26 week) | 0.552 | 1.344 [1.223, 1.464] | < 0.001 |
| LR 2 | INF B - Australia (present) | 0.627 | 0.655 [0.605, 0.705] | < 0.001 |
| LR 3 | INF B - U.S. (before 26 week) | 0.713 | 0.706 [0.578, 0.835] | < 0.001 |
|  | INF B - Australia (present) |  | 0.442 [0.383, 0.501] | < 0.001 |

Beta, Beta coefficient; CI, Confidence Interval; INF, Influenza; ILI, Influenza-like illness; LR, Linear regression; US, United States of America

All statistical analyses were performed using Python 3.6.2 (Python Software Foundation), and *p* < 0.05 was considered statistically significant.

### Prediction model

Forecasting seasonal influenza after 26 weeks was defined as forecasting influenza pattern after six months (26 weeks) with data available only at the current point (S2 Fig). Because of the time-consuming nature of vaccine production, vaccine strategy needs to be prepared at least six months in advance of the upcoming flu season [14]. Three seasons for 2015–2018 were selected to validate the prediction models for seasonal influenza patterns because the four seasonal surveillance data for Australia and Chile from 2011 to 2014 for the training set accounted for 50% of eight seasons in whole data. Moreover, influenza activity from the 41^th^ in 2018 to the 14^th^ in 2019 used external validation set. For example, the prediction model for the seasonal influenza pattern during 2015–2016 used the influenza surveillance data from the 40^th^ week in 2010 to the 40^th^ week in 2015 as the training set (S3 Fig).

The prediction model is a single-output model for seasonal influenza patterns after 26 weeks using the four historical values that indicate a month (4 weeks): we predict *Y*_t+26_ based on *X*_*t*_, *X*_*t*−1_, *X*_*t*−2_, and *X*_*t*−3_ [15].

Deep learning is a type of machine learning based on neural networks. The input layer receives an input signal, which moves to the next layer as a modified version of the input signal. It passes through several layers composed of multiple transformations, and last passes through the output layer as an output signal [16, 17]. The main algorithms for deep learning are deep neural networks (DNN), deep convolutional neural networks, deep belief networks, and recurrent neural networks. We selected DNN using the python library Keras (version 2.2.0) with TensorFlow (version 1.8.0) backend. The scikit–learn library (http://scikitlearn.org/) was used for data management and preprocessing. In this study, we used a three-layer DNN network with a 10% dropout rate for the overfitting problem. The models were optimized using the Adam optimizer with a loss function of mean squared error. Neuron activation functions were rectified linear units for the second layer. We selected 100 epochs and one batch size for the DNN model.

Prediction models of LR were calculated to forecast influenza surveillance after 26 weeks in the U.S. per each year. The influenza time series is characterized by an autocorrelation, so we also employed Auto Regressive Integrated Moving Average including exogenous variables (ARIMAX). The autocorrelation function and partial autocorrelation function is used to determine the autoregressive (AR) and moving average (MR) order. An ARIMA model is notated as ARIMA (*p, d, q*), where *p* indicates the AR order, *d* the differencing order and *q* the MA order [18]. Prediction models of LR and ARIMA were used the same input variables for deep learning in S1 and S2 Tables, respectively.

### Validation for the prediction model

The coefficient of determination, *R*^2^, is the proportion of the variance in the dependent variable that is predictable from the independent variables. The best possible score is 1.0, and it can be negative (because the model can be arbitrarily worse). Root-mean-square error (RMSE) was calculated using real values and predicted values for influenza activities in the validation set from the 41^th^ week in 2015 to 14^th^ week in 2019. The onset of influenza weeks for ILI is defined as the weighted percentage that exceeds the national baseline [19]. The peak amplitude and the peak timing are defined as the maximum value and that week in seasonal influenza week.

## Results

### Cross–correlation analysis

Table 1 shows the maximum correlation coefficient and time lag between the seasonal influenza outbreaks in the U.S. and the other countries. The correlation coefficients were calculated by cross-correlation between ILI in the U.S. and the number of positive influenza viruses worldwide. In Table 1, Australia had the highest correlation coefficient (0.896) with a −22 week time lag for the INF A. The −22 week time lag meant that the country’s seasonal influenza outbreak 22 weeks earlier was highly correlated with the seasonal influenza outbreak in the U.S.

In the analysis of the ILI and Total INF, the correlation coefficients of Australia were 0.855 with a −22 week lag and 0.892 with a −22 week lag, respectively. Chile had the third highest correlation coefficient, with a −28 week lag for ILI and INF A. However, the correlation coefficients of Australia for the INF B were 0.792 with a −25 week lag, which is much lower than that for the INF A. Moreover, the correlations for the INF B in such European countries as Norway, Croatia, the UK, and Sweden were higher than those of Australia in the Southern Hemisphere. We selected Australia and Chile to forecast ILI, Total INF and INF A after 26 weeks in the U.S., and Australia was selected for INF B.

Fig 2 (a) and (b), showing the INF A and INF B in the U.S. and Australia, shows that the values for Australia were shifted forward 22 weeks from the 40^th^ week in 2010 to the 40^th^ week in 2018. Interestingly, the waveforms for the U.S. and Australia are similar in peak timing and amplitude, and the dominant subtypes of the annual INF A are similar, except for 2014–2016. Fig 2 (c), showing the INF A in the U.S. and Chile, shows that the values for Chile was shifted forward 28 weeks, the waveforms also similar. Moreover, the dominant subtypes of the annual INF A are similar between the U.S. and Chile, except for 2016–2017. The subtype influenza activities in the U.S. were shown in S4–6 Fig. Therefore, influenza surveillance data in Australia and Chile could be valuable for predicting an influenza outbreak after 26 weeks in the U.S.

**Fig 2.**
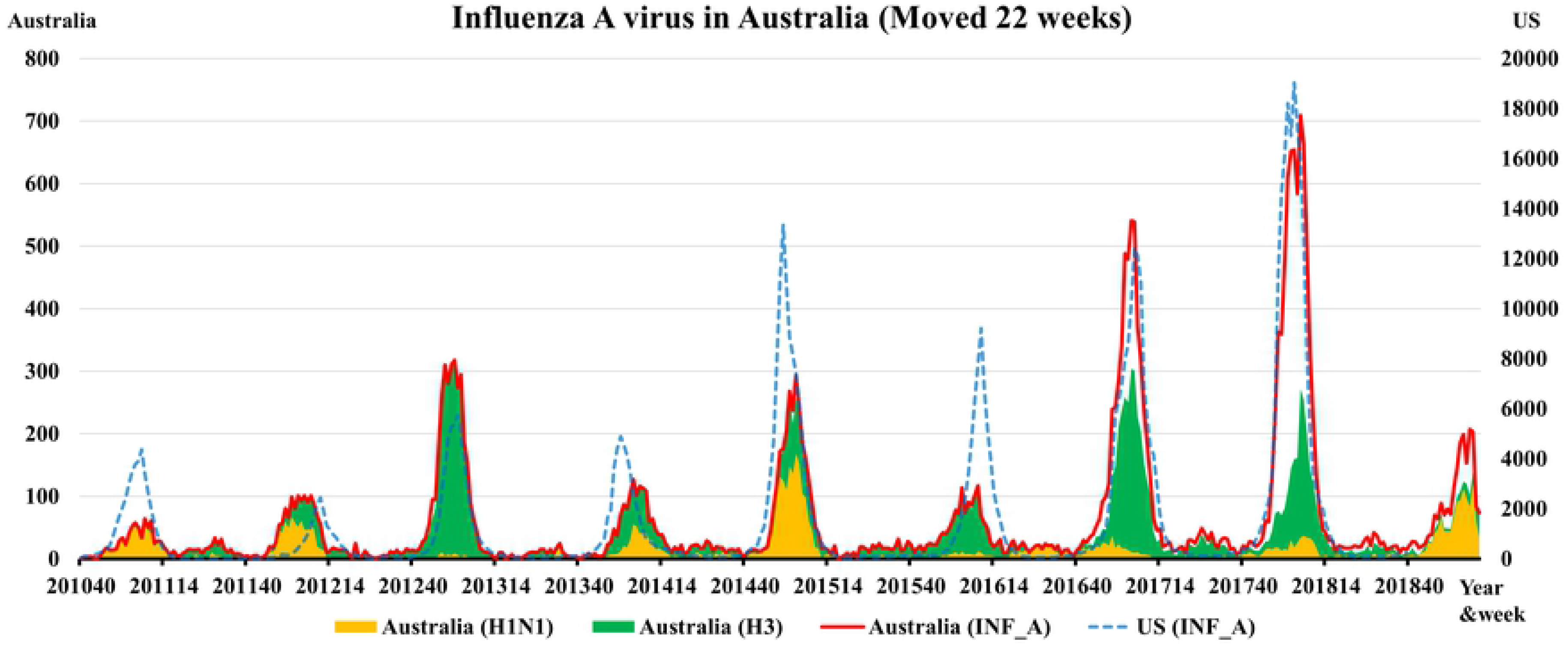
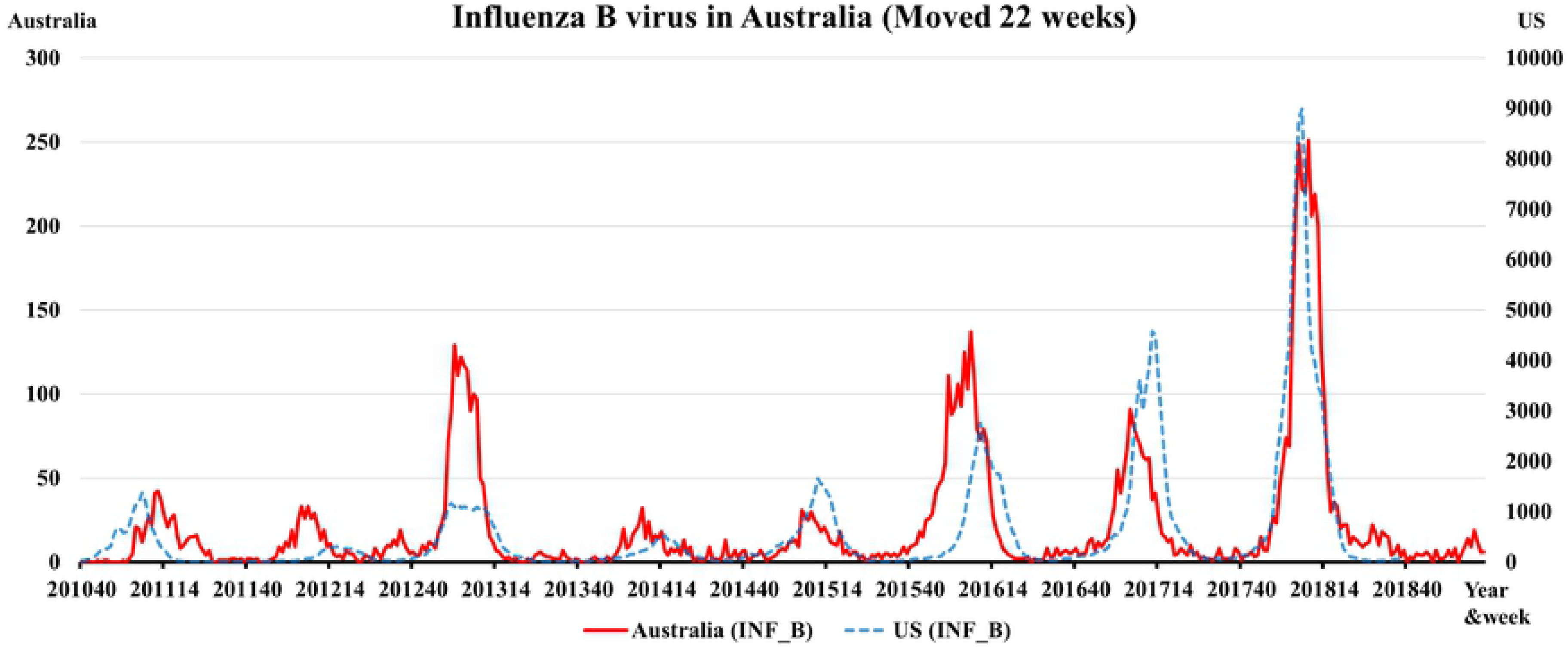
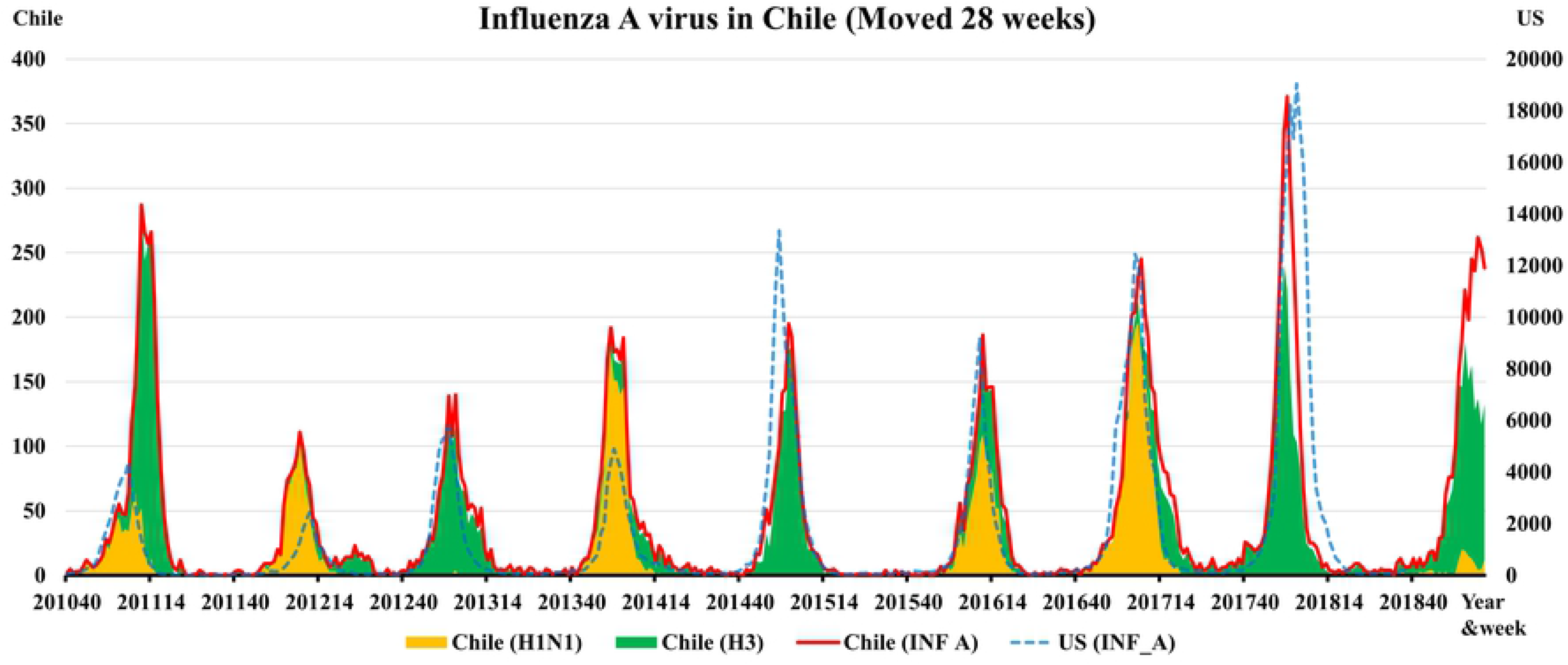
The surveillance data for influenza A (a) and B (b) viruses in the U.S. and Australia; the values for Australia were shifted forward 22 weeks in 2010–2018. The surveillance data for influenza A (c) viruses in the U.S. and Chile; the values for Australia were shifted forward 28 weeks. The blank part of graph is the number of influenza virus (not subtyped).

### Linear regression analysis

Table 2 shows results of LR for influenza after 26 weeks (ILI, Total INF, INF A, and INF B) in the U.S. using influenza activities for Australia, Chile, and previous season in the U.S. from 2010 to 2018. Results of LR demonstrated that adjusted R^2^ values for influenza surveillance in Australia and Chile at present were higher than those of previous season in the U.S. for forecasting those in the U.S. after 26 weeks for ILI, Total INF, and INF A. The adjusted R^2^ values for INF B in Australia also higher than those of previous season in the U.S. In LR 3, the influenza activities in Australia and Chile were significantly correlated with those after 26 weeks in the U.S. after adjusting for previous seasonal values in the U.S.

### Prediction models

The input variables of a prediction model for ILI after 26 weeks are ILI in the U.S at present and the Total INF in Australia and Chile at present, because of the absence of ILI data for other countries. The input variables for Total INF and INF A are Total INF and INF A, respectively, in Australia and Chile. The input variables for INF B are INF B in Australia.

Table 3 shows the performance of prediction models for seasonal influenza outbreaks in the U.S. after 26 weeks using previous season data, LR, ARIMAX, and DNN. The *R*^2^ score of DNN and ARIMAX for ILI, Total INF, INF A, and INF B showed better performance than those of previous season. *R*^2^ scores for the prediction models of DNN for ILI was 0.722. The all RMSE for forecast values and peak timing for the influenza week in DNN were less than those of previous season data. The RMSE for peak amplitude for ILI and INF B in DNN were less than those of previous season data, but those for Total INF and INF A in DNN were higher than those of previous season data.

**Table 3.** Performance of prediction models for seasonal influenza outbreaks after 26 weeks in the United States from 2015 to 2019.

| | | $R^2$ | RMSE<br>(forecast<br>values) | RMSE<br>(Peak<br>timing) | RMSE<br>(Peak<br>amplitude) |
| --- | --- | --- | --- | --- | --- |
| Influenza-like illness | Previous<br>season | 0.487 | 1.1 | 5.5 | 2.2 |
|  | LR | 0.328 | 1.2 | 1.2* | 3.1 |
|  | ARIMAX | 0.544 | 1.0 | 2.8 | 1.9* |
|  | DNN | 0.722* | 0.8* | 1.6 | 1.9 |
| Total Influenza viruses | Previous<br>season | 0.346 | 4802.8 | 5.8 | 6657.8* |
|  | LR | 0.589 | 3804.6 | 2.7 | 8021.7 |
|  | ARIMAX | 0.593 | 3786.3 | 1.7* | 6963.3 |
|  | DNN | 0.654* | 3490.0* | 1.9 | 7365.9 |
| Influenza A virus | Previous season | 0.289 | 4190.9 | 5.8 | 4901.0* |
|  | LR | 0.692* | 2759.8* | 1.9 | 5892.4 |
|  | ARIMAX | 0.635 | 3002.8 | 1.9 | 5456.4 |
|  | DNN | 0.678 | 2820.0 | 1.9* | 5704.8 |
| Influenza B virus | Previous season | -0.238 | 1880.8 | 3.7 | 5142.3 |
|  | LR | -1.436 | 2638.1 | 4.4 | 3736.0 |
|  | ARIMAX | 0.346* | 1367.4* | 2.2* | 2205.6 |
|  | DNN | 0.186 | 1524.7 | 3.5 | 1195.4* |

R<sup>2</sup>, Coefficient of determination; RMSE, Root-mean-square error; LR, Linear regression; ARIMAX, Auto Regressive Integrated Moving Average; DNN, Deep neural network.
\* Best value among previous season, LR, ARIMAX, and DNN.
Units for the RMSE (forecast values) are percentage of visits for influenza-like illness and number of total influenza, A, and B viruses.
Units for the RMSE (Peak timing) are week.
Units for the RMSE (peak amplitude) are percentage of visits for influenza-like illness and number of total influenza, A, and B viruses.

Fig 3 shows the prediction of DNN for ILI, Total INF, INF A, and INF B from the 41^th^ week in 2015 to the 14^th^ week in 2019 in the U.S. using the surveillance data from 26 weeks before in Australia and Chile. Moreover, the prediction and 95% confidence interval of ARIMAX for type specific influenza activities in the U.S. were shown in S7 Fig. The pattern of peak timing and amplitude of influenza activity for 2015–2016 season was different with those of 2012–2015 season, but our models forecast the later and less severe for influenza activity. Moreover, the predicted INF B activity for the 2018–2019 season was a better match than the increasing pattern for the 2013–2018 seasons. In Fig 3, the peak value of ILI for the U.S. in 2018–2019 could be less than that in 2017–2018. The onset and peak timing for the influenza week could be the 51^th^ week in 2018 and the 8^th^ week in 2019, which are later than those in 2017–2018.

**Fig 3.**
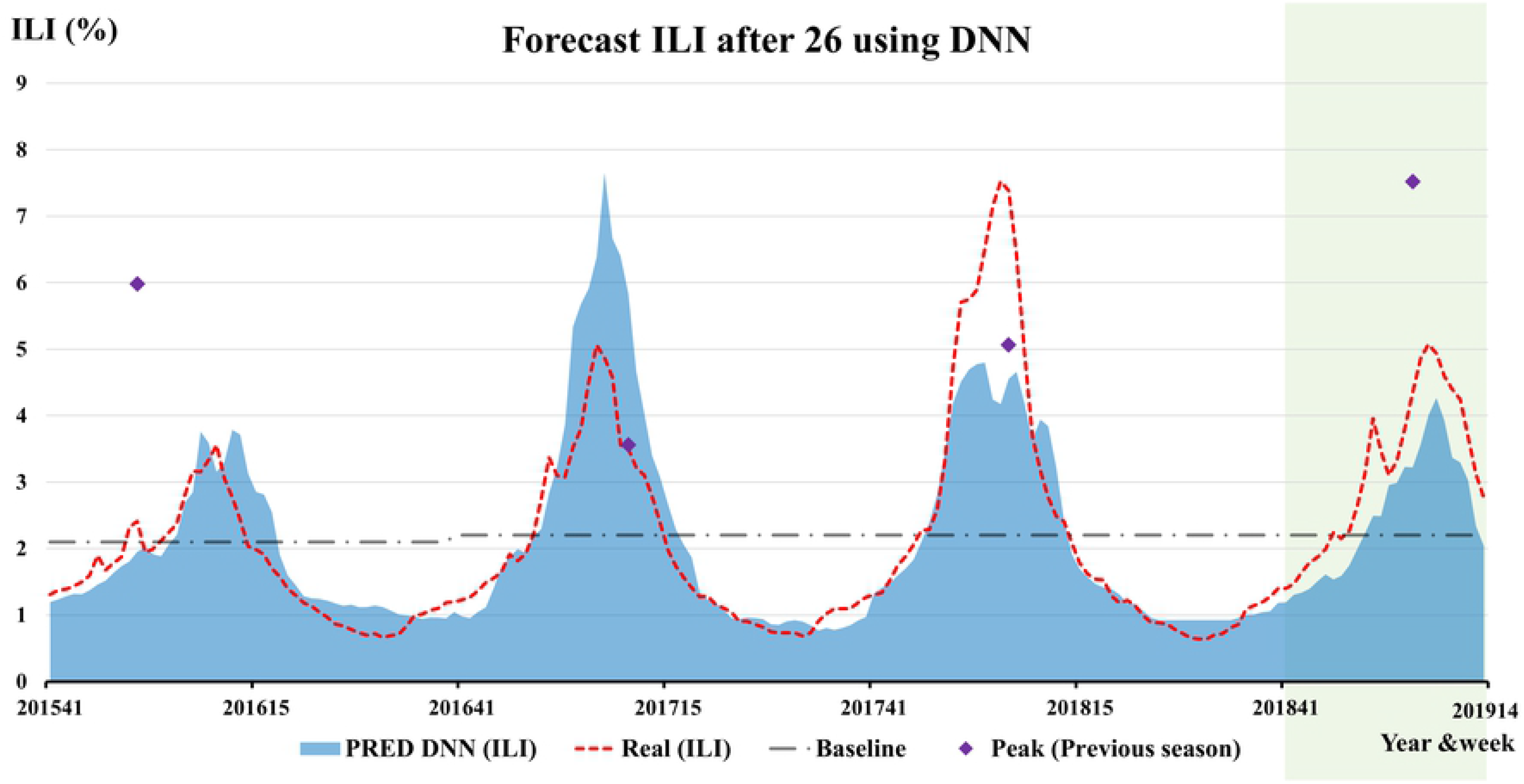
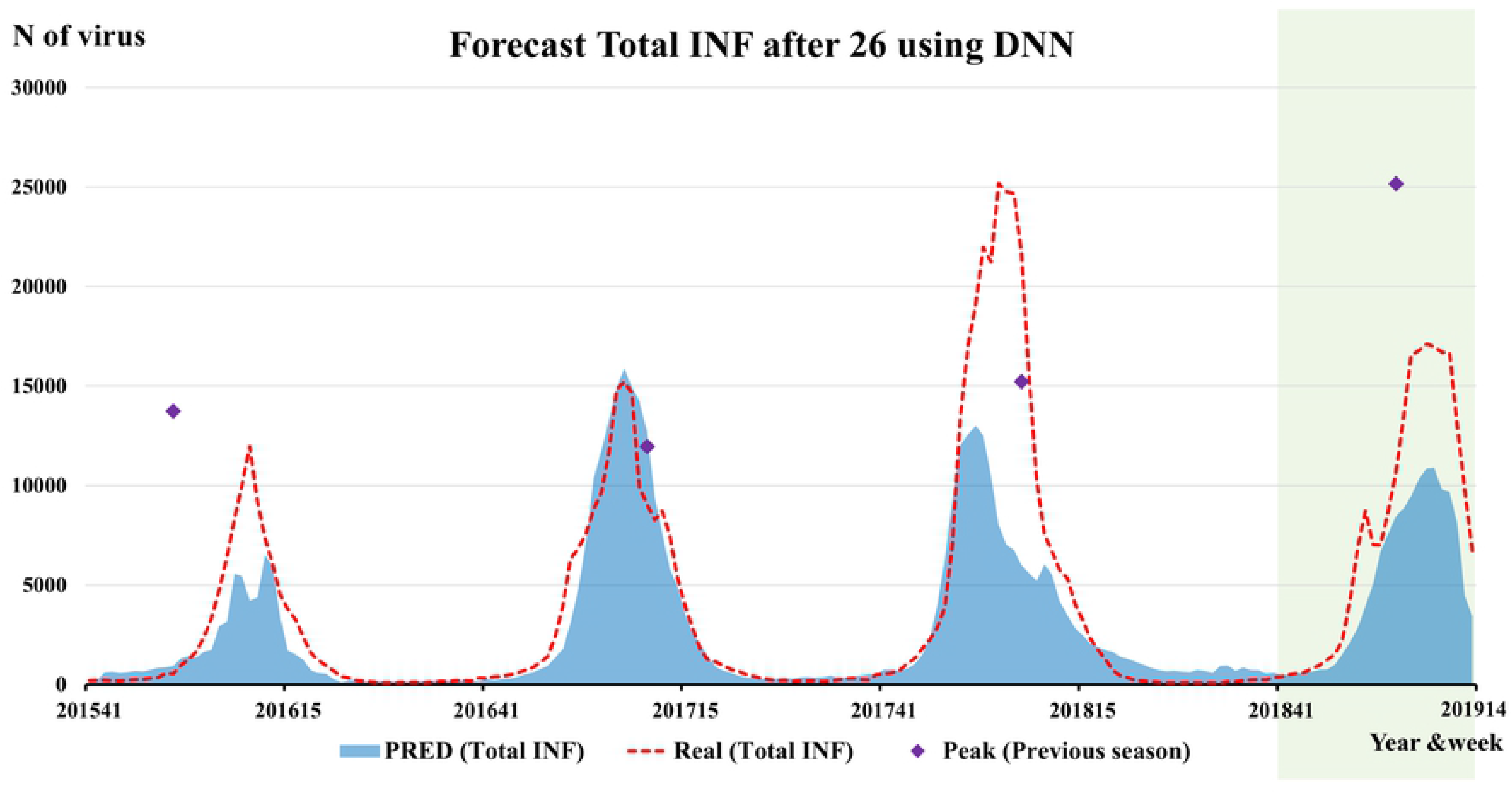
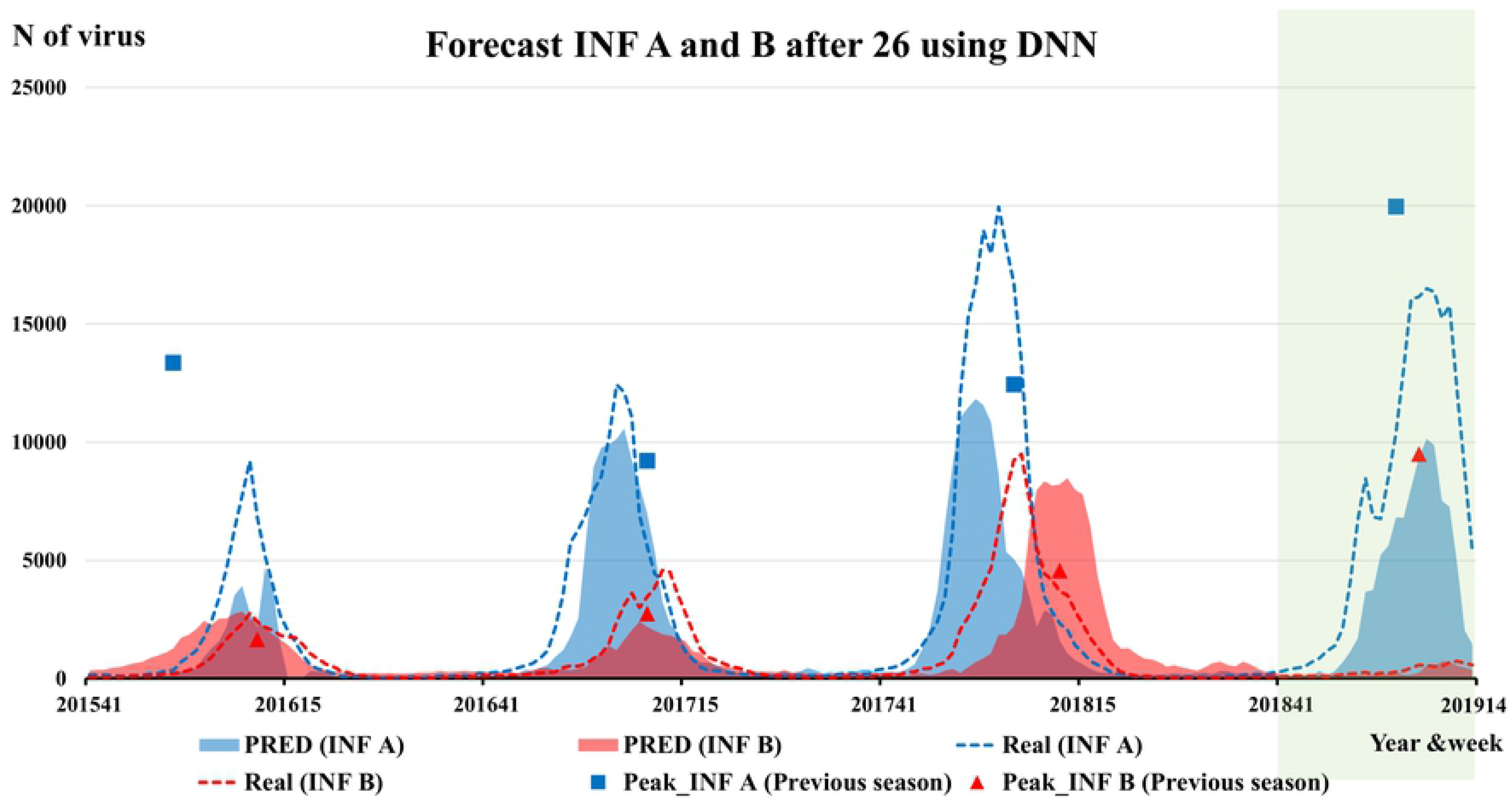
The prediction of DNN for ILI (a) and Total influenza (b), A and B viruses (c) from 2015 to 2019 in the U.S., using the surveillance data from 26 weeks before in Australia and Chile.

## Discussion

The aim of this study was to identify countries with seasonal patterns and influenza outbreaks that were similar to but preceded those of the United States. We used influenza surveillance of Australia and Chile to forecast for seasonal influenza after 26 weeks in the U.S. The seasonal influenza patterns in Australia before 22 weeks and Chile before 28 weeks showed a high correlation with those of the U.S. However, the correlation for the INF B between the U.S. and Australia was lower than that for the INF A. In Table 2, influenza surveillance in Australia and Chile at present were more useful than previous seasonal influenza in the U.S. for forecasting next seasonal influenza in the U.S. DNN models showed better performance for forecasting influenza activities after 26 weeks in the U.S than previous season data. The ARIMAX model also showed high performance which can be interpretable. Our prediction models forecast that the ILI for the U.S. in 2018–2019 may be later and less severe than that in 2017–2018.

The seasonal influenza patterns in the U.S. were highly correlated with those in Canada, Australia, Chile, and the United Kingdom. The correlation coefficients for these countries were higher than for their neighboring country or for other countries in the Northern Hemisphere, for example, Mexico (0.409 for Total INF), Spain (0.775), and Japan (0.599), which are not shown in Table 1. Moreover, correlated countries for the INF B were different from those for the INF A. The high correlation coefficients indicate similarities of peak timing, peak height, and onset timing for influenza outbreaks between two countries. For these reasons, the correlations for seasonal influenza between countries could be caused by characteristics of influenza, economic status, educational status, or access to health care as well as by seasonal environmental factors in the Northern Hemisphere [3, 20]. However, this study did not prove the causality of correlations for seasonal influenza between countries. Although we do not know the reasons, the patterns for seasonal influenza in the U.S., Australia, and Chile were similar, and influenza surveillance in Australia and Chile can be used to predict seasonal influenza outbreaks after 26 week in the U.S.

A study of Bedford *et al.* demonstrated that the less-frequent global movement of INF A/H1N1 and B viruses coincided with slower rates of antigenic evolution, lower ages of infection, and smaller, less-frequent epidemics than for A/H3N2 viruses [21]. Their study analyzed the correlation of peak timing for seasonal influenza between countries, and gave a time-series graph for influenza surveillance from 2000 to 2012 in the U.S. and Australia [21]. The time-series of virological characterizations for A/H3N2 in the U.S. and Australia were consistent with the time-series graph in our Fig 2(a). Bedford *et al.* suggested that differences in ages of infection could explain patterns of global circulation across a variety of human viruses [21].

Viboud *et al.* analyzed correlations for influenza epidemics from 1972 to 1997 in the U.S., France, and Australia [22]. In their study, France and the U.S. had a high correlation for influenza epidemics, but there was no significant correlation between the U.S. and Australia. In the scenario in which the influenza season in Australia was systematically six months in advance of that in the Northern Hemisphere, the median time lag between the peaks in Australia and in the United States was 27 weeks (range 14–39) [22]. Our study analyzed correlations for INF A and INF B, but the study of Viboud *et al.* did not [22].

The previous studies for prediction models for seasonal influenza have focused on social networking service data, search engine query data, and environmental factors [23–25]. These predictors are correlated with present influenza cases with a relatively short-term gap, of about one to four weeks. However, the influenza surveillance data in Australia had a time gap of 22 weeks from those in the U.S., which can help to establish a data-driven influenza vaccine strategy about six months ahead.

Kandula et al. analyzed whether forecasts targeted to predict influenza by type and subtype during 2003–2015 in the U.S. were more or less accurate than forecasts targeted to predict Total INF using four compartmental models [26]. They found that forecasts separated by type/subtype generally produced more accurate predictions and, when summed, produced more accurate predictions of Total INF [26]. Our prediction models for type-specific influenza is valuable as well as those for ILI, which could provide an important, richer picture of influenza activity.

To our best knowledge, this is the first study to demonstrate the relevance of influenza surveillance in Australia and Chile for predicting influenza cases in the U.S. The performance of our prediction model could be improved by integrating influenza surveillance data in other countries, internet search data, and self-reported data.

Our study has several limitations. For the ILI study, we could not use ILI data in other countries, instead had to use data for Total INF, because collecting ILI data separately for the 164 countries was difficult. There were not enough data on other potential covariates to show relationships between influenza outbreaks in Australia, Chile, and the U.S., such as the standard of the medical facilities, economic level, and medical records of the influenza virus. We did not show uncertainty intervals for prediction results of DNN models, but the 95% confidence interval for prediction of ARIMAX were shown in S7 Fig. Furthermore, we included only data from laboratory-confirmed cases, which may underestimate the true incidence of influenza in the population [27]. Further research with inter-US geographic scales for US Health and Human Services regions is warranted to analyze detail scales to find matched country with similar pattern of influenza surveillance.

## Conclusions

Our study forecasts the 2018–2019 seasonal influenza after 26 weeks in the U.S. using the 2018 seasonal influenza in Australia and Chile. Although the predicted values may be different from the actual values, the correlation for seasonal influenza between the two countries could be used to decide on influenza vaccine strategy about six months ahead in the U.S. Our prediction model allows to estimate peak timing, peak intensity, and type-specific influenza activities for next season at 40^th^ week.

## Supporting information

**S1 Fig.** The example of the cross-correlation analysis.

**S2 Fig.** The explanation for forecasting seasonal influenza after 26 weeks.

**S3 Fig.** The explanation for training set at 40^th^ week and output for forecasting seasonal influenza after 26 weeks.

**S4 Fig.** The surveillance data for influenza-like illness, Total influenza, A, and B viruses in the U.S. in 2010–2018.

**S5 Fig.** The surveillance data for H1N1 and H3 viruses of influenza A type in the U.S. in 2010–2018.

**S6 Fig.** The surveillance data for Victoria and Yamagata viruses of influenza B type in the U.S. in 2010–2018.

**S7 Fig.** The prediction and 95% confidence interval of Auto Regressive Integrated Moving Average for ILI (a) and Total influenza (b), A (c) and B viruses (d) from 2015 to 2019 in the U.S.

**S1 Table.** Linear regression models for influenza surveillance after 26 weeks in the U.S.

**S2 Table.** Auto Regressive Integrated Moving Average including exogenous variables for influenza surveillance after 26 weeks in the U.S.

## Acknowledgments

This work was supported by a National Research Council of Science and Technology grant from the Korean government (MSIP) (no. CRC–16–01–KRICT). We also acknowledge the National Influenza Centers (NICs) of the World Health Organization’s Global Influenza Surveillance and Response System (GISRS).

## Data Availability

ILI data in the U.S. are available from the Fluview database of the Centers for Disease Control and Prevention in the U.S. The agency’s website https://gis.cdc.gov/grasp/fluview/fluportaldashboard.html

Influenza surveillance data are available from the FluNet database of the WHO Global Influenza Surveillance Network http://apps.who.int/flumart/Default?ReportNo=12

## Author Contributions

Conceptualization: SBC, IA.

Data curation: SBC, JK.

Formal analysis: SBC, IA.

Methodology: SBC, JK.

Supervision: IA.

Writing – original draft: SBC.

Writing – review & editing: SBC, JK, IA

